# Bayesian semantic surprise based on different types of regularities predicts the N400 and P600 brain signals

**DOI:** 10.1101/2024.03.13.584794

**Authors:** Alice Hodapp, Alma Lindborg, Milena Rabovsky

## Abstract

The brain’s remarkable ability to extract patterns from sequences of events has been demonstrated across cognitive domains and is a central assumption of predictive processing theories. While predictions shape language processing at the level of meaning, little is known about the underlying learning mechanism. Here, we investigated how continuous statistical inference in a semantic sequence influences the neural response. 60 participants were presented with a semantic oddball-like roving paradigm, consisting of sequences of nouns from different semantic categories. Unknown to the participants, the overall sequence contained an additional manipulation of transition probability between categories. Two Bayesian sequential learner models that captured different aspects of probabilistic learning were used to derive theoretical surprise levels for each trial and investigate online probabilistic semantic learning. The N400 ERP component was primarily modulated by increased probability with repeated exposure to the categories throughout the experiment, which essentially represents repetition suppression. This N400 repetition suppression likely prevented sizeable influences of more complex predictions such as those based on transition probability, as any incoming information was already continuously active in semantic memory. In contrast, the P600 was associated with semantic surprise in a transition probability model over recent observations, possibly indicating a working memory update in response to violations of these conditional dependencies. The results support probabilistic predictive processing of semantic information and demonstrate that continuous update of distinct statistics differentially influences language related ERPs.

## 1. Introduction

Many events in our daily lives are not very surprising because they follow a certain pattern or structure. These patterns allow for some predictability regarding sequences of events, from weather patterns and social behavior down to music and language. But how can we make these predictions? One proposal within the general framework of predictive processing is that the brain is constantly estimating and adapting the underlying generative model of the environment and is making predictions based on these estimates (Bayesian framework: e.g., Friston, 2005, 2010; Rao & Ballard, 1999; but see also e.g., Elman, 1990; McClelland, 1994; Rogers & McClelland, 2008).

The ability to track these regularities is a crucial mechanism in human cognition, spanning sound (e.g., Saffran et al., 1999), visual (e.g., Fiser & Aslin, 2001, 2005; Monroy et al., 2019), tactile (e.g., Conway & Christiansen, 2005) and motor sequences (e.g., Baldwin et al., 2008) even in non-human primates (e.g., Hauser et al., 2001; Newport et al., 2004) and songbirds (e.g., Abe & Watanabe, 2011). One open debate is what type of regularity is inferred by the brain. In electroencephalography (EEG), various event-related potential (ERP) components seem to be modulated by the frequency of exposure to a stimulus. In these experiments, rare stimuli (deviants) are more surprising and elicit a larger neural response. Deviants have been operationalized in different ways: they can be infrequently presented across the experiment (e.g., stimulus A is presented more often than B across an experiment; Garrido et al., 2009, 2013; Lecaignard et al., 2015; Näätänen et al., 2007) or result from a local repetition of a stimulus even if there is no overall frequency bias in the sequence (e.g., a local pattern of AAAB; Kolossa et al., 2013; Squires et al., 1976; Ulanovsky et al., 2004). However, neural responses in such sequences can also be explained by conditional probabilities, for which expectations depend on an event that already occurred. Specifically, items that are surprising based on transition probabilities (probability of current stimulus given the previous stimulus) elicit a larger response, oftentimes reflected in the same ERPs previously suggested to be modulated by frequency-based statistics (Gijsen et al., 2021; Higashi et al., 2017; Koelsch et al., 2016; Maheu et al., 2019; Marcovitch & Lewkowicz, 2009; Meyniel et al., 2016; Mittag et al., 2016). Moreover, some studies suggested that frequency of exposure and transition probability might predict the ERP at different latencies and integrate information across different timescales (Maheu et al., 2019; Pesnot Lerousseau & Schön, 2021).

Transition probabilities have also been demonstrated to be essential for language acquisition and processing. Seminal work by Saffran, Aslin, and Newport (1996) demonstrated that infants can track transition probabilities between syllables in an artificial language, which has been interpreted as the mechanism supporting word segmentation in continuous speech for language acquisition (e.g., Estes et al., 2007; Mirman et al., 2008). Literature proposes that statistical learning effects extend to the phonotactic (Chambers et al., 2003), orthographic (Pacton et al., 2001), and syntactic (Gomez et al., 2000; Saffran, 2001; Ullman, 2004) level. In this study, we want to explore statistical learning at the level of meaning – arguably the central aspect of language processing.

There is a lot of evidence in the sentence processing literature that the brain can use patterns in language to make semantic predictions. This has often been investigated by violating semantic expectations and recording participants’ EEG. The N400 is characterized as a centro-parietal negative going brain potential that peaks around 300-500 ms after word onset (Kutas & Hillyard, 1980). It is gradually reduced with higher predictability of an incoming stimulus from the preceding context (Kutas & Federmeier, 2011; Kutas & Hillyard, 1980, 1984) or semantic overlap with the predicted continuation (Federmeier & Kutas, 1999). Based on this sensitivity to predictability, the N400 is often interpreted as an index of unpredicted semantic information (Bornkessel-Schlesewsky & Schlesewsky, 2019; Federmeier et al., 2007; Kuperberg, 2016; Kuperberg et al., 2020; Lindborg et al., 2023; Rabovsky et al., 2018). However, the N400 is not the only ERP that has been discussed in the context of predictive processing of semantic information. A subsequent posterior positivity between approximately 600 and 800 ms is also often observed after a semantically anomalous continuation. This positivity is referred to as a semantic P600 effect (Brothers et al., 2020; Hoeks et al., 2004; Kim & Osterhout, 2005; Kuperberg et al., 2003; Münte et al., 1998) to differentiate it from the P600 associated with syntactic ambiguities and violations (Friederici et al., 1993, 1996; Hagoort et al., 1993; Münte et al., 1998; Osterhout & Mobley, 1995). These P600 findings are not necessarily contradicting or representative of two separate ERP components, but might rather reflect a more general signal of error monitoring in language (Ryskin et al., 2021; van de Meerendonk et al., 2009, 2010).

The current study investigated which statistics are inferred by the brain to make semantic predictions and how this learning modulates language related ERPs. One interpretation of the N400 is that its amplitude reflects semantic prediction error and/ or Bayesian surprise, which has previously been implemented in algorithmic-level neural network models whose N400 correlates approximate these computational level concepts (Fitz & Chang, 2019; Rabovsky et al., 2018; Rabovsky & McRae, 2014). In line with this proposal, Lindborg et al. (2023) recently modeled N400 amplitudes as Bayesian surprise at the level of meaning in an oddball-like paradigm consisting of a continuous sequence of nouns that belonged to different semantic categories. Semantic sequences are characterized by the well-replicated N400 priming effect, meaning that the N400 is reduced when words from the same category are presented successively and is increased when the category switches, resulting in a semantic mismatch effect (Bentin et al., 1985; Holcomb, 1988; Holcomb & Neville, 1990; Rugg, 1985). Semantic priming in sequences can be explained in terms of statistical learning and probabilistic predictive processing, as any new information will update an internal probabilistic representation (i.e., generative model) to reduce future prediction errors. If one or multiple items that share semantic features are processed, the internal model adapts its estimates, and these features will be less surprising when presented again, resulting in reduced neural activity (see Rabovsky et al., 2018; Rabovsky & McRae, 2014 for discussion). Lindborg et al. (2023) showed that semantic surprise in a model continuously adapting its probability estimates for the semantic categories based on exposure during recent trials predicted trial-by-trial amplitude changes in the N400. These results not only support the assumption that the brain is trying to infer and constantly adapt the current generative model but also propose recent exposure as a possible measure that is used to make predictions.

Here, to investigate the influence of semantic surprise beyond recent exposure, we employed a similar semantic statistical learning paradigm as Lindborg et al. (2023) with an additional transition probability manipulation between categories, meaning that each semantic category followed one specific semantic category more frequently than the other semantic categories. We devised two models to capture different aspects of probabilistic learning. Both models were Bayesian sequential learner models, from which we derived a trial-by-trial semantic surprise measure based on a categorical representation of the stimuli, in which each incoming word belongs to one of five semantic categories.

The first model estimates the probability of the category of the next stimulus based on (recent) exposure, following Lindborg et al. (2023). To recap, this model has previously captured the classic N400 oddball effect at the level of meaning – e.g., the presentation of a musical instrument (e.g., cello) after a sequence of vegetables (e.g., tomato, carrot, cucumber) is more surprising than the presentation of another vegetable. This model is referred to as category exposure (CE) model. In addition, we devised a more complex model that tracks not only how frequently a category has recently occurred but also how surprising a category is specifically after a particular preceding category (e.g., how probable is a musical instrument after a vegetable as opposed to after a tool). This model is referred to as transition probability (TP) model. These transition probabilities between categories would not influence surprise in the category exposure model. The recency of previous experience that the model considers is based on the timescale of integration that is optimized during model fitting (see Figure 1).

**Figure 1.**
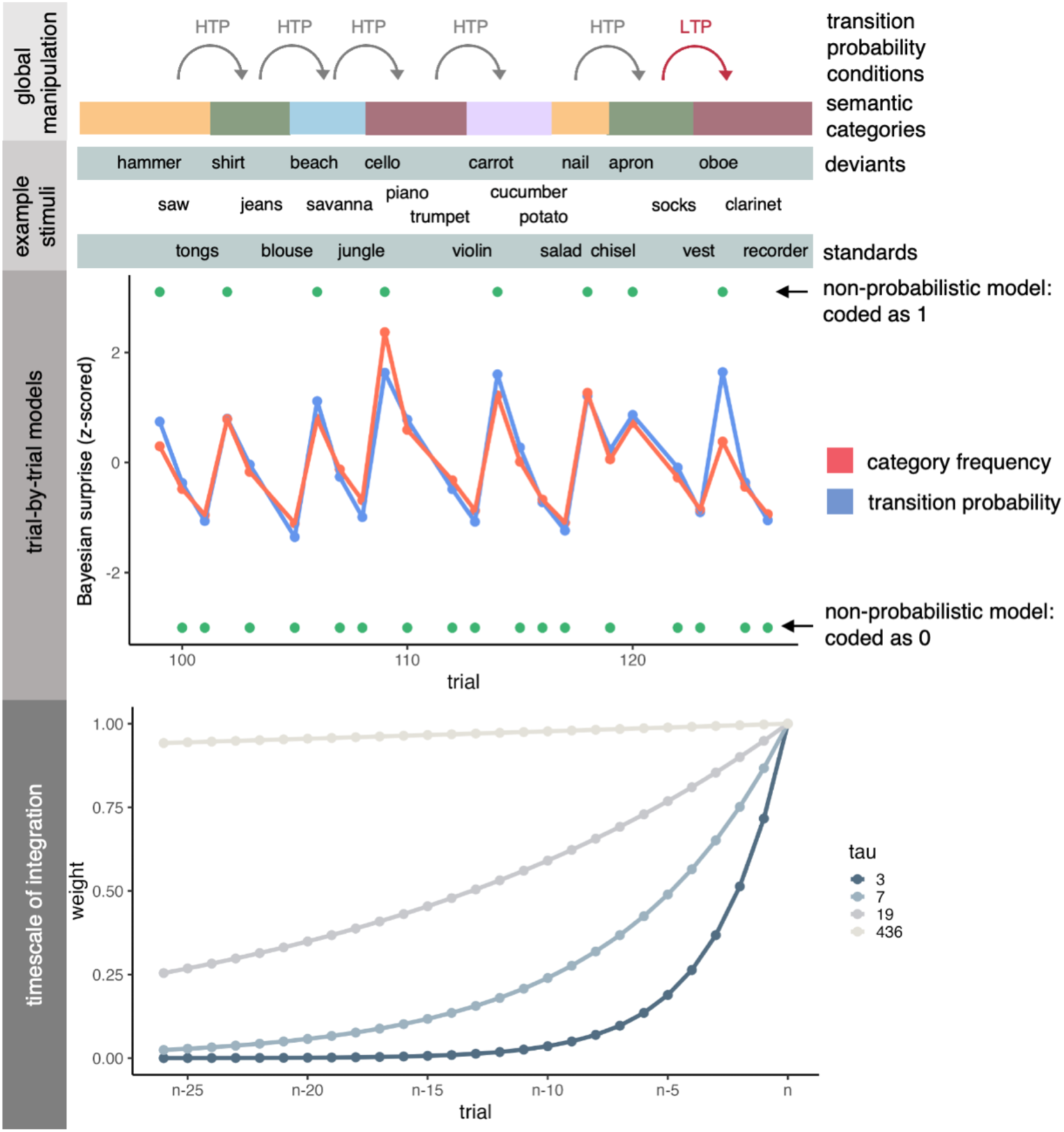
First panel: On a global level, experimental items belonged to one out of five semantic categories, illustrated by the colored blocks. The transition from one category to another was either 85% (high transition probability, HTP) or 5% (low transition probability, LTP) across the entire experiment, which is the basis for the condition assignment in the test phase. Second panel: Nouns from the categories formed sequences of varying lengths that were presented item-by-item in a continuous oddball-like roving paradigm. The first item of a category sequence is labeled as deviant, and the last item as standard for the ERP analysis. Third panel: Example output for all model types. For both probabilistic models (category exposure, CE, and transition probability, TP) the semantic surprise decreases within the category sequences. Green dots indicate the coding of a non-probabilistic baseline model which labeled deviants as 1 and all other trials as 0. Probabilistic models are plotted with *1*=10. Non-word trials are missing from the plot. Fourth panel: The value of *τ* determines the memory weighting function. Trials with a memory weight of 1 are fully included in the learner’s probability estimate, whereas a trial with a weight of 0 is fully excluded. This means that with a lower value of *τ*, the model includes only relatively recent trials, whereas with a larger *τ*, more trials are included. We refer to the number of trials included as timescale of integration.

Both models simulate the behavior of a participant in an experiment who implicitly compares each new incoming word to their current beliefs about the semantic context and adjusts their beliefs according to the new input. If incoming information has a high likelihood based on the model estimate (based on previous exposure or transitions) it results in small surprise and minimal adjustment to the learner’s beliefs. Information with low likelihood, however, will prompt the learner to reconsider the probabilities of the respective semantic categories, which is reflected in larger semantic surprise and stronger belief adjustment.

By relating the model derived theoretical semantic surprise levels to the EEG response, conclusions about the mechanism reflected in the N400 and P600 can be drawn. First, if modulations of the N400 and/ or P600 in our noun sequence are driven by a probabilistic representation of previous exposure (i.e., semantic priming effects), we expect semantic surprise in the CE model to significantly predict these amplitudes. If modulations for the N400 and/ or P600 are driven by statistical learning of more complex features, i.e., differences in transition probability between categories, we expect semantic surprise in the TP model to significantly predict these components. Second, if the two components represent predictions based on different time scales, we expect the optimal integration timescale for the models to differ when fitted to the respective components.

Previous findings from sentence processing show clear effects of sentence context beyond overlap in recently processed semantic features (i.e., semantic priming) on the N400 and P600 ERP components (Kutas & Federmeier, 2011; Van Petten & Luka, 2012). Transition probabilities as implemented in the current abstract sequence paradigm can be seen as a very basic approximation to the use of context cues to anticipate upcoming semantic information. We therefore expected semantic surprise based on not just recent exposure but also transition probabilities to modulate both N400 and P600 amplitudes.

Overall, the current study included two approaches towards understanding which statistics influence semantic predictions. In addition to the two theoretical surprise measures described above, which were calculated trial-by-trial to account for participants’ EEG modulations while they were continuously exposed to the to-be-learned statistics, we also recorded ERPs in a subsequent test phase. In this test phase, we compared the event-related response based on the type of transitions between trials: within category transitions (e.g., a tool after a tool), between category transitions (e.g., a tool after a musical instrument), which were further divided into high probability and low probability transitions according to the participants’ experience during learning (see Figure 1 and Figure 2), and novel transitions (e.g., a tool after a land animal, which was never presented during continuous learning). These two approaches are complementary in that they capture both online probabilistic semantic learning and the influence of the learned statistics on the semantic mismatch response.

**Figure 2.**
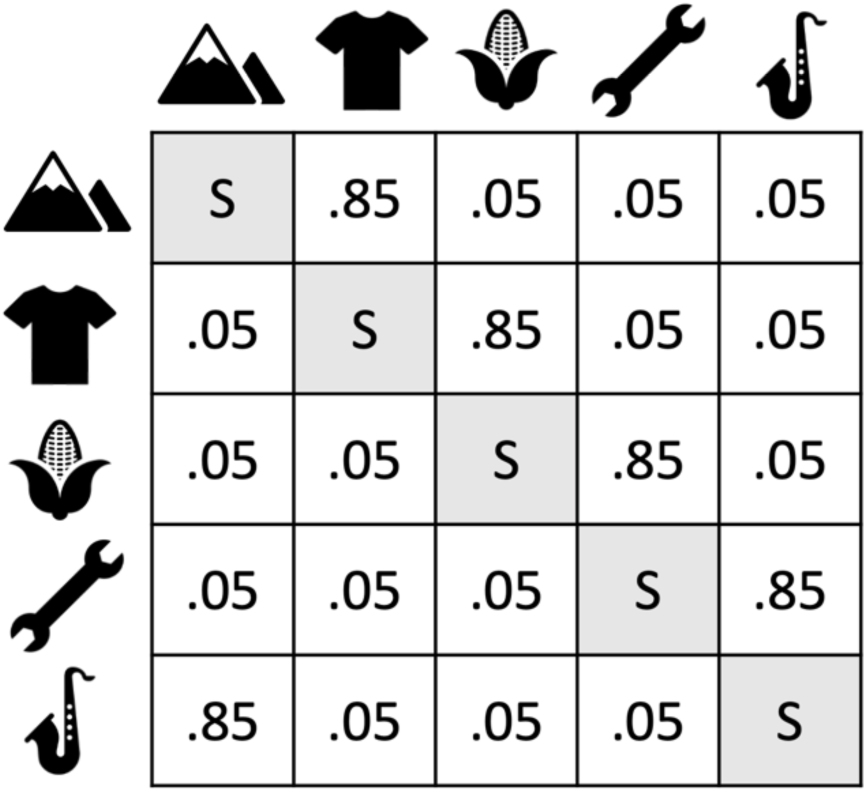
Global transition probabilities between the categories. Within category transitions on the diagonal are labeled as standard (S) for the ERP test phase analysis. The probabilities are conditional on a category change i.e., the probabilities of category as deviant given the previous standard. The category assignments are counterbalanced across participants. Transitions of 85% are labeled HTP (high transition probability deviant) and transitions of 5% are labeled LTP (low transition probability deviant) for the ERP test phase analysis. Please note that this does not reflect the trial-by-trial probabilities estimated by the Bayesian sequential learner.

**Figure 3.**
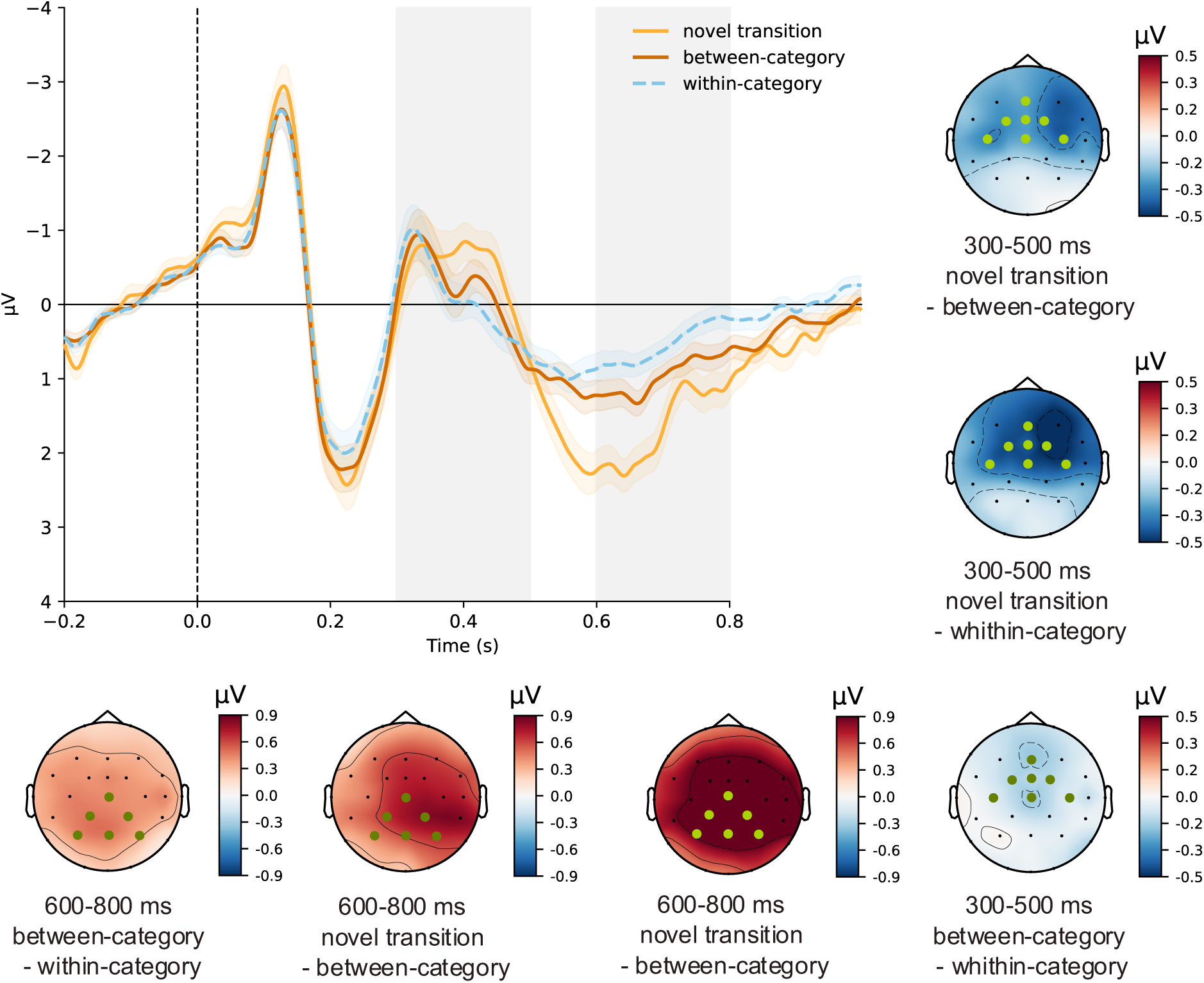
Test phase ERP results. Grand-average waveforms at fronto-central ROI with regression-based baseline correction based on the 200 ms before stimulus onset. The 300-500 and 600-800 ms time window is shaded in gray. Error bands indicate the SEM. The ROI for the respective statistical analysis is marked with green dots.

## 2. Methods

### 2.1. Participants

The experiment is part of a research project whose protocols were approved by the Ethics Committee of the German Psychological Association (DGPs; MR102018). All participants gave written informed consent before the experimental session in accordance with the Code of Ethics of the World Medical Association (Declaration of Helsinki). 60 volunteers (17 male) between the ages of 18 and 38 (mean age: 23.43) participated in the experiment. Three participants had to be replaced due to excessive artifacts. Participants were compensated by either course credits or money. All were native German speakers, right-handed (Edinburgh Handedness Inventory; Oldfield, 1971), had normal or corrected-to-normal vision, and none reported a history of neurological or psychiatric disorders.

### 2.2. Stimuli

Stimuli consisted of 50 German nouns belonging to 5 semantic categories. Each of the categories (landscapes, vegetables, clothes, tools, and musical instruments) contained 10 nouns. The stimuli were restricted to have no more than one commonly used meaning and to be reasonably well-known. Further, the stimuli were selected to have high within-category and low between-category semantic similarity and were thus optimized for a categorical model (see below). It was verified in separate one-way factorial ANOVAs that the categories did not significantly differ on any of the following control variables: absolute type frequency (*F*(4) = 0.032, *p* = 0.86) and number of orthographic neighbors according to Coltheart (1977) (*F*(4) = 0.74, *p* = 0.39), both extracted from dlexDB German language corpus (Heister et al., 2011), as well as the number of letters per noun (*F*(4) = 0.92, *p* = 0.34). As a control task, to ensure that participants paid attention, participants were reacting to non-words. These non-words were all pronounceable and had no orthographic neighbors in German.

### 2.3. Procedure

#### 2.3.1. Main sequence: Oddball-like roving paradigm

To study semantic statistical learning and single-trial surprise signals, stimuli were presented in an oddball-like roving paradigm (see also Lindborg et al., 2023). In the task, a sequence of nouns from the same semantic category (e.g., different landscapes) was presented to the participants one by one on a screen. The sequence was followed immediately by a series of nouns from a different category (e.g., clothes) with the categories repeating to create a continuous sequence. Comparable to a classic oddball paradigm, the last noun belonging to a category can be regarded as a *standard* (expected) and the first noun of the next category as a *deviant* (unexpected), with the advantage that deviants and standards can be perfectly counterbalanced across the experimental session. See Figure 1 for an illustration of the paradigm.

The main sequence of the experiment consisted of a total of 450 category trains (each category is presented 90 times), which consisted of 2 to 4 nouns. These category trains were constructed such that the probability of a given noun occurring at a specific position within a category sequence and the number of repetitions across items was balanced. Overall, participants saw 1450 experimental items (+3 dummy items that comprised the first category, since this first category doesn’t have a transition, see below). Unknown to the participants, we additionally manipulated the probability of one category being followed by another. Each category had one high transition probability category (HTP), which is a transition that happened in 85% of all category changes occurring after that preceding category. Each of the other three categories followed in 5% of all category changes after that preceding category (low transition category: LTP). These probabilities reflect the probability of a specific category as deviant given the previously presented standard. This assignment of transition probabilities was also counterbalanced across participants. Outside of the counterbalancing constraints, the presentation of the nouns within categories and category changes were randomized within participants. The probabilities in Figure 2 illustrate the manipulation of the deviants and are the basis of the ERP conditions used in the test phase analysis (see next section). They are conditioned on a category change; hence, they do not represent the trial-by-trial transition probabilities estimated by the TP model. In the learner model diagonal elements will dominate, as within-category transitions are more common than category switches due to the roving design.

To ensure that participants were paying attention, they were instructed to react with a button press to non-words that were interspersed in the stimuli train. The 120 non-words occurred every 8-24 words and never directly before or after the last noun in a category (i.e., a standard).

#### 2.3.2. Test phase: New semantic categories

To analyze the effect of the additional transition probability manipulation (Figure 2) on the semantic mismatch effect in a classic grand-average ERP analysis, the experiment contained a subsequent test phase in which the design of the oddball-like roving paradigm changed slightly. If global statistics can be used to make predictions, participants should have a representation of these statistics after being exposed to the main sequence. This test phase of the experiment contained 180 presentations of trains from the five known categories (same categories and same pool of nouns as in the main sequence) and the presentation of 50 noun trains from completely new categories. These new categories were not analyzed but enabled us to split the deviants (first item after category change) into three conditions, based on the category that was presented in the previous trial: 100 high transition probability deviants (HTP according to the main sequence of the experiment), 30 low transition probability deviants (LTP according to the main sequence of the experiment) and 50 novel transition deviants (after a new category, so that no transition can be predicted based on previous experience). Importantly, the deviants themselves were all known items from the five known semantic categories. Sequence length and noun position within the category sequence were balanced.

### 2.4. Data acquisition and pre-processing

EEG data was recorded at 32 active Ag/AgCl electrodes (actiCHamp, Brain Products) mounted in an elastic electrode cap based on the international 10-20 system (Jasper, 1958), while eye movements were monitored with bipolar electrodes at the outer canthi of the eyes. EEG and EOG were recorded with a sampling rate of 1000 Hz and electrode impedances were kept below 5 kΩ. Preprocessing was performed using EEGlab and ERPlab toolboxes (Delorme & Makeig, 2004; Lopez-Calderon & Luck, 2014) and custom scripts. Noisy channels were identified in 26 participants (only one channel for each of the participants) and interpolated using spline interpolation. Data were then re-referenced against the average of the left and right mastoid, filtered with a 0.1 Hz high pass (two-pass Butterworth with a 12 dB/oct roll-off) and a 30 Hz low pass filter (two-pass Butterworth with a 24 dB/oct roll-off) and downsampled to 500 Hz. Eye movement, blinks, heart, and channel noise components were corrected using independent components analysis (ICA) and the IClabel plugin (Pion-Tonachini et al., 2019). The IC weights were transferred from continuous data filtered at 1 Hz to the correctly filtered data as described above. IClabel then automatically removed all components that were classified as the respective category with a probability > 30% (average of 4.95 ± 1.64 components per participant). The corrected signal was segmented from −200 to 1000 ms time-locked to stimulus onset. Segments with values exceeding ±75 µV were automatically marked for rejection. This left an average of 1430 ± 29 trials for analysis of the main sequence and an average of 777 ± 16 trials for analysis of the test phase.

### 2.5. ERP analysis

For the statistical analysis, mean amplitudes were extracted in a priori determined time windows of 300-500 ms (N400) and 600-800 ms (P600) after stimulus onset and averaged across electrodes within a region of interest. In line with other results from mass repetition studies (Debruille & Renoult, 2009; Lindborg et al., 2023), the N400 effect for this experiment was more central than its typical centro-parietal distribution (e.g. Kuperberg et al., 2020). We therefore adjusted the region of interest (ROI) to include FC1, FCz, FC2, C3, Cz, C4, CP1 and CP2. For the test phase (and after more repetitions) the distribution was even more fronto-central, so the ROI included Fz, FC1, FCz, FC2, C3, Cz, and C4 (however, main results did not change with a more central ROI). The mean amplitudes in the 600-800 ms time window were extracted in a centro-parietal ROI (Cz, CP1, CP2, P3, Pz, P4), as typical for the P600.

We then performed linear-mixed effect model (LMM) analyses using the package lme4 (Bates et al., 2014) as implemented in R (R Core Team, 2022). Following the recommendations of Barr et al. (2013), we tried to fit the maximal random effect structure as justified by the design and reduced its complexity successively until the model converged. The 200 ms baseline period was included in the statistical models as a covariate (Alday, 2019). The significance of fixed effects was determined via likelihood ratio tests, comparing the fit of the model to that of a model with the same random effects structure without the respective fixed effect.

For the data from the test phase, we first compared within-category transitions (standards only), between-category transitions (combining HTP and LTP deviants), and novel transitions (deviant occurring after a new category was presented) with treatment coding (within-category as a baseline). Note that novel transitions are of course also between-categories but have *additionally* not been experienced before. For completeness, we added difference coding, comparing between-category deviants to within-category and then novel transitions to between-category transitions. This was then followed-up by splitting the between-category transition condition into a comparison between LTP and HTP trials (sum coding: -0.5, 0.5). For the main sequence of the experiment, deviants and standards were compared regarding an overall semantic mismatch effect using sum coding (-05, 0.5).

### 2.6. Trial-by-trial analysis

For a single trial analysis of the main sequence, we derived the regressors from simple Bayesian sequential learners that were then fitted to the single trial EEG data. The trial-by-trial learning that is reflected in the model assumes that the probabilistic expectation for a given incoming word is different for each presentation depending on what was learned in the unique combination of previous trials. Two model types were used: a model learning the probability of each semantic category based on previous exposure in the sequence (category exposure model) and a model learning the transition probabilities between semantic categories in the sequence (transition probability model). It is important to note, that while the *overall* frequency of exposure to each of the semantic categories in the experiment was kept constant, the sequence was still characterized by *local* differences in exposure frequency (see implementation of memory constraints below; also see Figure 1) that could influence trial-by-trial neural responses, first because the order of the semantic categories was randomized and second because recent observations are often from the same category due to the oddball like roving design (see Lindborg et al., 2023 for a similar design). Regressors were used across the whole post-stimulus time window and all electrodes to identify EEG signals that are modulated by the different statistics modeled here.

### 2.7. Semantic surprise models

The Bayesian sequential learners were based on a categorical representation of the stimuli, in which each noun encountered by the learner falls into one of the five semantic categories. For simplicity, we first consider the Bayesian category exposure model (CE).

The CE model was implemented as a Dirichlet-Categorial model, that keeps count of the categories observed to derive the best estimate of their probability (Gijsen et al., 2021; Lindborg et al., 2023). The likelihood of observing a stimulus of the semantic category given the current beliefs follows a categorical distribution:

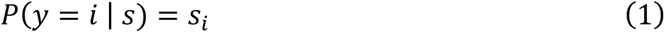

where *s* = (*s*_1_, …, *s*_5_), ∑ *s_i_* = 1 are the observer’s current beliefs of the probability of observing a stimulus from each semantic category. The beliefs *s* follow a Dirichlet distribution with parameters (*α*_1_, …, *α* _5_), which all have an initial value of 1 (reflecting equal probability of observing any category) and are updated by calculating the posterior distribution *p*(*s* | *y_t_*) after observing the stimulus *y_t_*. Applying this approach to each stimulus sequentially, we get the following belief distribution at time *t*:

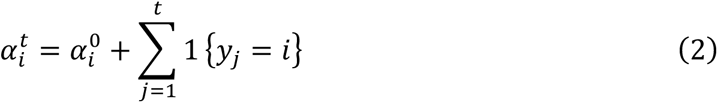

To account for biological memory constraints, the models have a limited timescale of integration (leaky integration). In our model, this was implemented as an exponential decay on the previous trials in the calculation of the beliefs *α^t^*, defined by

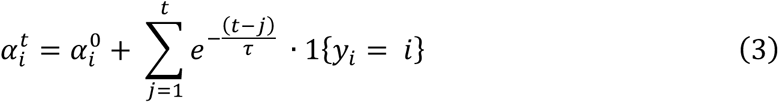

where the parameter *τ* controls the length of the window of integration (small *τ* yielding a shorter window and large *τ* yielding a long window). For a more detailed description and full derivation of these results, see Lindborg et al. (2023). The estimation of transition probabilities (TP model) was implemented by counting types of transitions instead of exposure to categories. The transition probability matrix describes how likely each category is, given its preceding context i.e., the previous category in the sequence. Hence, the model infers five sets of probability vectors (one for each category) each describing the model’s current beliefs of the probability of observing a stimulus from each semantic category next. Note that this model estimates the full transition vector for a given category (i.e., transitions within the same category take the highest value). These vectors are different from the probabilities conditional on a category change (i.e., deviants) in Figure 2 which illustrates the experimental design and conditions for the test phase ERP analysis.

For both Bayesian models described above, prediction error (PE; based on predictive surprise) and Bayesian surprise (BS) were implemented as possible surprise read-outs. In short, predictive surprise is defined as the predictive likelihood of the item perceived (Shannon, 1948), implemented as the negative logarithm of the posterior predictive distribution *p*(*y*_*t*_|*s*_*t*_). If the posterior probability for the observation *y*_*t*_ is low, it will cause a large prediction error. Bayesian surprise (Itti & Baldi, 2009) is defined as the Kullback-Leibler (KL) divergence between the model’s beliefs before (prior) and after (posterior) the observation of the current item (Modirshanechi et al., 2021). An example of how BS varies over a sequence of stimuli in the oddball-like roving paradigm is displayed in Figure 1. For a more detailed description and full derivation of Bayesian surprise at trial *t*, see Lindborg et al. (2023).

Semantic surprise was calculated over semantic and not lexical features, which is an important distinction from the classic linguistic notion of surprisal which refers to surprise over individual lexical items (Frank et al., 2015; Levy, 2008). Semantic surprise was calculated separately on each participant’s stimulus sequence and z-scored. In line with Lindborg et al. (2023), we use a category switch measure as a non-probabilistic baseline model, which assumes maximal semantic surprise when the current item is from a different semantic category compared to the previous item, and no semantic surprise otherwise. The deterministic model therefore codes deviants as 1 and all other trials as 0.

### 2.8. Model comparison

Each combination of model type (category frequency and transition probability) and surprise readout function (Bayesian surprise and prediction error) resulted in unique regressors for each stimulus sequence, which were fitted to baselined single-trial, event-related electrode data. EEG data and continuous regressors were z-scored. The timescale of integration *τ* was optimized for each subject and model combination. As previous research reported a very local timescale of integration for a category exposure model in a semantic oddball task (*τ* = 3; Lindborg et al., 2023), we implemented each integer up to 10 plus an additional 55 values of *τ* in a logarithmically spaced vector to cover the entire interval from 1 to perfect integration (*τ* = ∞). Previous research focused specifically on the N400 time window and ROI (Lindborg et al, 2023), however, the current analysis of the overall mismatch effect (deviant versus standard) and the regression ERP approach (see below) revealed an additional late time window of interest. This raises the possibility that the brain estimates statistics computed across multiple timescales in parallel (e.g., Bernacchia et al., 2011; Kiebel et al., 2008; Meder et al., 2017; Ulanovsky et al., 2004), which may be reflected in the two effects of interest. Critically, this includes the possibility that one statistic can explain both effects based on different timescales of information integration as implemented via different values of *τ* (see Figure 1). Therefore, regressions were estimated at each time point and electrode and then optimized (the only free parameter was *τ*) in two time windows of interest (300-500 ms and 600-800 ms) (see Maheu et al., 2019 for a similar approach regarding τ). These time windows of interest correspond to the N400 and P600 ERP components and are thus in line with previous ERP literature on predictive language processing.

Surprise readouts (BS and PE) within models were compared using Bayesian model selection, which treats models as random effects in the population and can successfully deal with population heterogeneity and outliers (Stephan et al., 2009). For the analysis, we approximated the marginal likelihood (or model evidence) of each model (CE and TP) and surprise measure (BS and PE), by using the Bayesian information criterion. The random-effect analysis was performed as described in Stephan et al. (2009) and Rigoux et al. (2014), and as implemented in the VBA toolbox (Daunizeau et al. 2014). The analysis returns the expected model frequency and the exceedance probability *χπ*, i.e., the probability for each model to be more frequent than any other model in the general population. We report protected exceedance probabilities that are corrected for the possibility that observed differences in model evidence (across participants) are due to chance (Rigoux et al., 2014).

### 2.9. Regression ERPs

Using the surprise read-out that was selected via Bayesian model comparison (see last section), participant-specific surprise estimates from both models were regressed against baselined single-trial EEG data using time-resolved linear regression, for which the estimated *β* coefficients of each predictor (CE and TP model) can be visualized as ERP-like plots depicting how semantic surprise modulates neural activity across time (regression evoked response: rERP; Smith & Kutas, 2015). For this analysis, EEG data was low pass filtered with a 10 Hz filter (for a similar filter see also Heilbron et al., 2022; Kern et al., 2022) and downsampled to 250 Hz. EEG data and continuous regressors were z-scored. Analysis was performed using custom Python code and the MNE rERP implementation (Gramfort et al., 2013; Smith & Kutas, 2015). Please see e.g., Heilbron et al. (2022), Kern et al. (2022) and Visalli et al. (2021) for similar regression-based approaches within the predictive processing literature.

Statistical testing was conducted a cluster-based permutation test (Maris & Oostenveld, 2007) with threshold-free cluster enhancement (TFCE; Mensen & Khatami, 2013) to correct for multiple comparisons (start: 0, step: 0.2). Mass-univariate testing was performed using one-sample t-tests with hat variance adjustment (σ = 1e^-3^; Ridgway et al., 2012). By-channel time courses were plotted as heatmaps indicating data points exceeding the 5% threshold (settings and plots based on Sassenhagen, 2019).

## 3. Results

### 3.1. Test phase: Semantic mismatch analysis

Figure 3 plots the grand average ERPs for the test phase of the experiment, grouping standards and deviants in terms of their learned transition probabilities. We first compared within-category transitions (standards), between-category transitions (combining LTP and HTP deviants), and novel transitions (deviants after a new category). In the 300-500 ms time window, novel transitions elicited a more negative N400 than within-category (*β*=-0.4, *SE*=0.12, *χ*^2^=11.76, *p*<0.001). Novel transitions were also had a more negative N400 than and between-category transitions (*β*=-0.28, *SE*=0.13, *χ*^2^=4.68, *p*=0.03) with a frontal N400 distribution (as typical for mass repetition studies; Debruille & Renoult, 2009; Lindborg et al., 2023). There was no significant difference for between-category and within-category transitions (*β*=-0.14, *SE*=0.09, *χ*^2^=2.73, *p*=0.09).

The 600-800 ms time window revealed a graded P600 effect: between-category elicited a significantly more positive ERP than within-category transitions (*β*=0.23, *SE*=0.08, *χ*^2^=8.51, *p*=0.003), while novel transitions were more positive than between-category transitions (*β*=0.60, *SE*=0.12, *χ*^2^=26.50, *p*<0.001). Each of these comparisons revealed a centro-parietal distribution, that most resembled the P600 ERP component.

A follow-up analysis split the deviants into transitions that were categorized as LTP (low transition probability) and HTP (high transition probability) based on the manipulation of transition probabilities in the main sequence of the experiment (Figure 2). There was no significant difference between these two types of transitions in the earlier N400 (*β*=0.17, *SE*=0.18, *χ*^2^=0.89, *p*=0.35) or later P600 time window (*β*=-0.04, *SE*=0.03, *χ*^2^=2.45, *p*=0.12). The grand-average ERP plot can be seen in Figure 4.

**Figure 4.**
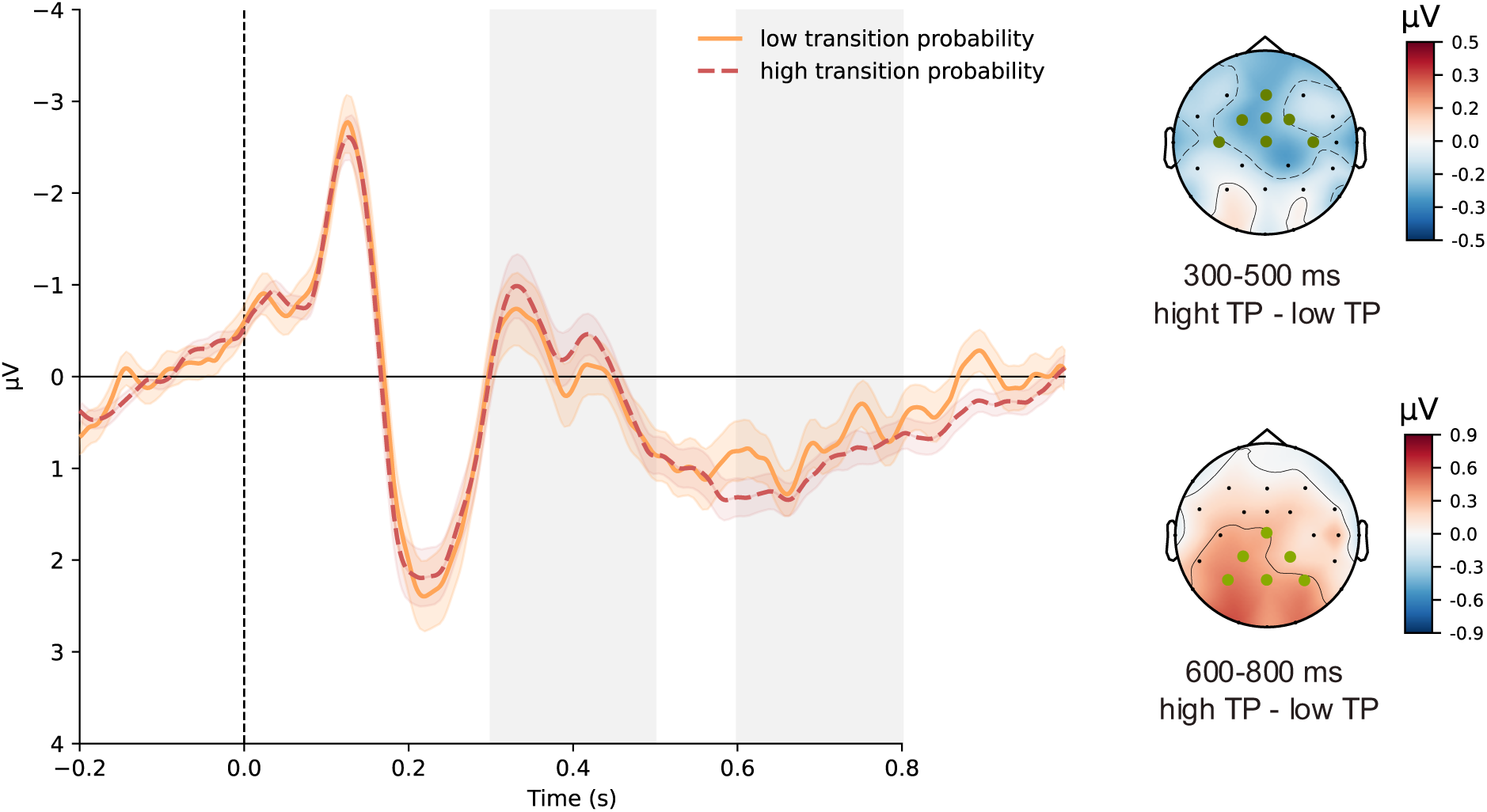
Test phase ERP results on transition probabilities. Grand-average waveforms at fronto-central ROI with regression-based baseline correction based on the 200 ms before stimulus onset. The 300-500 and 600-800 ms time window is shaded in gray. Error bands indicate the SEM. The ROI for the respective statistical analysis is marked with green dots.

### 3.2. Main sequence: Semantic mismatch analysis

Figure 5 depicts the standard (within category transition) and deviant (between category transition) comparison for the main sequence of the experiment. Deviants elicited a statistically more negative ERP compared to standards in the 300-500ms time window (*β*=-0.28, *SE*=0.06, *χ*^2^=15.4, *p*<0.001). In the 600-800ms time window deviants were characterized by a more positive response than standards (*β*=0.12, *SE*=0.04, *χ*^2^=5.67, *p*=0.017).

### 3.3. Main sequence: Bayesian sequential learner results

Bayesian model selection (BMS, as detailed above) was used to assess the relative plausibility of different surprise measures (Bayesian surprise versus prediction error) and a baseline model via posterior model probabilities and exceedance probabilities. Within the category exposure and transition probability model, the model estimating Bayesian surprise performed best, regardless of the time window to which parameters were fitted (300-500 ms and 600-800 ms). Additionally, both probabilistic learner models (CE and TP) continuously had a higher probability than the deterministic category switch null model which did not predict trial-by-trial changes in amplitude. In the earlier N400 time window, Bayesian surprise was deemed the best with high certainty for both the model learning the probability of categories based on recent exposure (p(MCF(BS)|y) = 0.987; *φ,*=1) and the model learning the categories’ transition probabilities (p(MTP(BS)|y) = 0.909; *φ*=0.94). When the model parameter was fit to the later P600 time window, the data revealed a similar pattern. For both category exposure (p(MCF(BS)|y) = 0.986; *φ*=1) and transition probability (p(MCF(BS)|y) = 0.934; *φ*=0.87), the model implementing Bayesian surprise had the highest posterior probability and protected exceedance probability. Therefore, all subsequent analyses reported were performed using Bayesian surprise estimates.

Results from the regression ERP and TFCE analysis revealed that recent category exposure and transition probability of semantic categories modulated the ERP in distinct time windows. Bayesian surprise from a category exposure model modulated the early and mid-latency neural response (0-500 ms), whereas Bayesian surprise in the transition probability model seemed to modulate late responses after 500 ms. This pattern was visible regardless of the time window to which the models’ time integration windows (*τ* parameter) were fitted.

When fit specifically to the 300-500 ms time window, Bayesian surprise in a category exposure model negatively modulated the neural response across most electrodes at around 100, 200, and most prominently around 400 ms with a fronto-central distribution (Figure 6 A and B). The best-fitting parameter values suggest a slow global timescale of integration for the category exposure learner in this time window, with a *τ* of 436 as the best-fitting value across participants (that is a half-life of ∼302 observations, Figure 6 E). Using the single-subject peaks, *τ* was found to significantly differ from perfect integration (one sample t-test: *p*<0.001). Coefficients for transition probability surprise from the same regression model showed no significant modulation of the ERP signal before 500ms, but a positive modulation for the late ERP signal.

When fitting the time window of integration specifically to the late 600-800ms time window, transition probability Bayesian surprise led to a more positive ERP in the time window after 500 ms across central electrodes, lateralized to the right (Figure 7 A and B). The best-fitting parameter values suggest a local timescale of integration for the transition probability learner fitted to this time window, with a *τ* of 7 as the best-fitting parameter across participants (that is a half-life of ∼5 observations, Figure 7 E). Using the single-subject peaks, *τ* was found to significantly differ from perfect integration (*p*<0.001) as well as from *τ*=1 (*p*=0.001). In the respective regression model (containing model estimates optimized for the 600-800ms time window) surprise in a category exposure model did not significantly modulate the ERP at any electrode or time point after 500ms.

Even though the TFCE analysis revealed no influences of Bayesian surprise based on transition probabilities before 500 ms (Figure 7C and Figure S2C in the supplement), as can be seen in Figure 7A (and Figure S2A in the supplement), the *ß* values from the grand average rERP seem to differ from zero not only in the P600 time window but also in the N400 time window. Because we had clear hypotheses concerning an influence of transition probability surprise also on N400 amplitudes, we performed a post-hoc test specifically on the N400 time window (300-500ms) in the depicted centro-parietal ROI (which corresponds to a very typical N400 ROI) with the above obtained by-participant *τ*. The linear-mixed model analysis indeed revealed a highly significant influence of transition probability surprise (*β* = *-0.26, SE*=0.001, *χ*^2^=830, *p* < .001) in addition to category exposure surprise. Of course, given the non-significant TFCE analysis in this time window, this significant post-hoc test is to be interpreted with care, and replication seems required.

For a visualization of the TP coefficient from the regression model fit to the 300-500 ms time window and the CE coefficient from the regression fit to the 600-800 ms window (i.e., the coefficients that showed a significant modulation *outside* the window to which the parameters were fitted) which were reported above, see figures in the supplementary material. Note that while the parameter *τ* was fitted to distinct time windows to account for the possibility that one model might explain effects at multiple latencies but with different timescales of integration, this was not the case and the results in the supplement lead to the same conclusions as the findings which are visualized in the figures in the main text. This is because the overall pattern for the best τ parameters by model type stayed consistent across both time windows of interest.

## 4. Discussion

In this study, we investigated how statistical learning at different levels of complexity influences EEG signatures. To this end, we devised an oddball-like semantic roving paradigm, in which nouns from different semantic categories are presented in sequences. The experiment tested for potential learning effects on processing in a test phase and additionally analyzed whether probabilistic models tracking previous exposure to semantic categories and transition probabilities between semantic categories can predict the EEG response in a meaningful way. While early and mid-latency brain responses were primarily modulated by surprise in a model estimating probabilities based on previous exposure to the semantic categories over a long timescale (global integration), late brain waves were modulated by surprise in a transition probability model reflecting recent observations (local integration).

The paradigm consisted of a noun sequence in which the last word of one semantic category can be considered a “standard” and the first word of the next semantic category can be considered a “deviant”. The EEG recorded during sequence processing was characterized by a semantic mismatch response (i.e., the difference between standards and deviants) in the N400 and P600 time window. The N400 mismatch response is in line with well-replicated results from semantic priming experiments, in which single words that were preceded by a semantically related word elicit a smaller N400 amplitude (Bentin et al., 1985; Holcomb, 1988; Holcomb & Neville, 1990; Rugg, 1985). This effect has also previously been found in semantic oddball sequences (Lindborg et al., 2023). While a P600 effect for semantic mismatch is less often reported, a late positive response has been described for some paradigms (Holcomb, 1988) and seems to be present in a previous semantic oddball experiment (but not analyzed: Lindborg et al., 2023).

Mismatch ERPs in auditory or somatosensory sequence processing have been suggested to reflect probabilistic Bayesian learning (Gijsen et al., 2021; Grundei et al., 2023; Kolossa et al., 2013; Lieder, Daunizeau, et al., 2013; Lieder, Stephan, et al., 2013; Mars et al., 2008; Squires et al., 1976) and first attempts have been made to map this assumption to the N400 in language processing (Lindborg et al., 2023; Delaney-Busch et al., 2019, but note that this latter paper refers to rational adaptation and see also Nieuwland, 2021). The current results support this assumption since trial-by-trial EEG signals for the sequence were better explained by probabilistic Bayesian models than a deterministic baseline model, suggesting that ERP responses elicited by semantic sequences are context-dependent and reflective of trial-by-trial statistical learning rather than a deterministic response to a pattern violation.

We also found that Bayesian surprise, i.e., the amount of update to the model’s beliefs caused by a stimulus, explained the data better than an alternative measure of prediction error (improbability of an event under the current beliefs), regardless of the model type (for a comprehensive overview of different measures of surprise, see Modirshanechi et al., 2021), which is in line with some theoretical accounts (see Lindborg et al., 2023; Rabovsky et al., 2018 for work on the N400 and model update). However, it is important to point out that the aim of this comparison between Bayesian surprise and prediction error was mainly to select a measure for further analysis, since both have previously been used in the relevant literature, and not to make claims on the exact computation performed by the brain. Without additional experimental manipulation, events inducing prediction error usually also induce a strong change in beliefs, and the two measures are therefore highly correlated.

Model update also depends on certainty which makes a paradigm with a varying stimulus quality, volatile environment, or instructions explicitly requiring update on only some of the trials (see O’Reilly et al., 2013; Visalli et al., 2021), a desirable future perspective to differentiate between Bayesian surprise and prediction error. A central finding of this study was that by using Bayesian surprise to model trial-by-trial neural responses, successive brain waves could be related to different types of statistical inferences and timescales of integration: early and mid-latency responses were primarily modulated by probability estimates based on previous exposure to semantic categories over a long timescale, while late responses were modulated by transition probability learning over a shorter timescale. This might suggest a difference in the mechanism underlying early and late neural signals that is supported by previous ERP findings and modeling work mostly in auditory processing. For example, Todorovic & de Lange (2012) showed that stimulus repetition attenuates early auditory responses (repetition suppression), while more complex stimulus expectation influences the following intermediate stages of processing. Using Bayesian sequential learner models of item frequency and transition probabilities, Maheu et al. (2019) demonstrated that successive brainwaves in auditory sequence processing are best explained by statistics of varying complexity and timescale of integration.

The exact timescale of integration for the results reported here is characterized by high uncertainty (a large variability is often found in literature, e.g., Gijsen et al., 2021; Maheu et al., 2019; Musiolek et al., 2019) and direct comparison to findings from paradigms outside language processing is difficult, due to the cofound between time and number of items (Pegado et al., 2010) and differences in complexity (Ma et al., 2014). However, the current results of an overall pattern of global integration for surprise in a category exposure model and a local integration for transition probabilities are in line with results on auditory processing described by Maheu et al. (2019).

In sum, our results from a semantic oddball paradigm suggest that the brain infers statistics on the level of meaning, that this is a probabilistic process, and that there seems to be a dissociation between stimulus repetition and more complex violated expectations. In the following, we will successively address the two main patterns regarding the estimated statistics as revealed by our ERP and sequential learner model results and discuss their implications and relation to other findings.

### 4.1. Bayesian surprise based on category exposure modulates early and mid-latency brain responses

When analyzing ERPs after the exposure to the main sequence in a subsequent test phase, there was no evidence for a significant difference between within-category transitions (standards) and between-category transitions (deviants) in N400 amplitudes. The absence of this semantic mismatch effect, which was present for the main sequence (Figure 5), can likely be explained by the mass repetition of the semantic categories that could have reduced N400 amplitudes across the experiment, diminishing the mismatch effect towards the test phase. However, novel transitions lead to a larger N400 amplitude. These novel transitions should be influenced by repetition suppression to the same extent as the other conditions since they consisted of the same five semantic categories and differed only with respect to the previous (novel) category. The ERP differences must therefore reflect a different mechanism. A tentative explanation could be that the previous processing of a novel category and/or a completely new transition indicated a change point in the environment, increasing uncertainty and with it the learning rate and attention (O’Reilly, 2013; Yu & Dayan, 2005).

**Figure 5.**
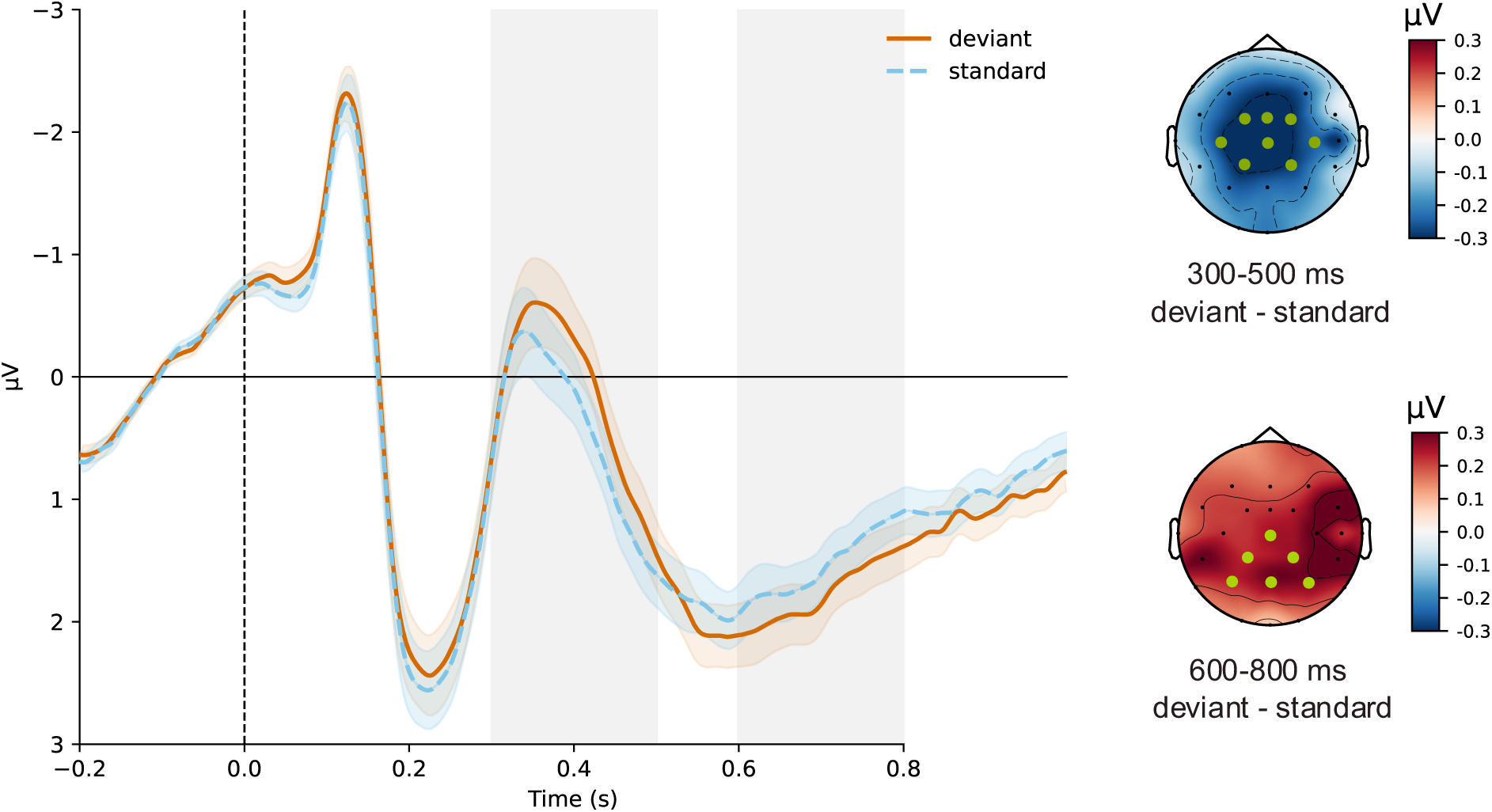
Main sequence ERP results. Grand-average waveforms at central ROI with regression-based baseline correction based on the 200 ms before stimulus onset. The 300-500 and 600-800 ms time window is shaded in gray. Error bands indicate the SEM. The ROI for the respective statistical analysis is marked with green dots.

**Figure 6.**
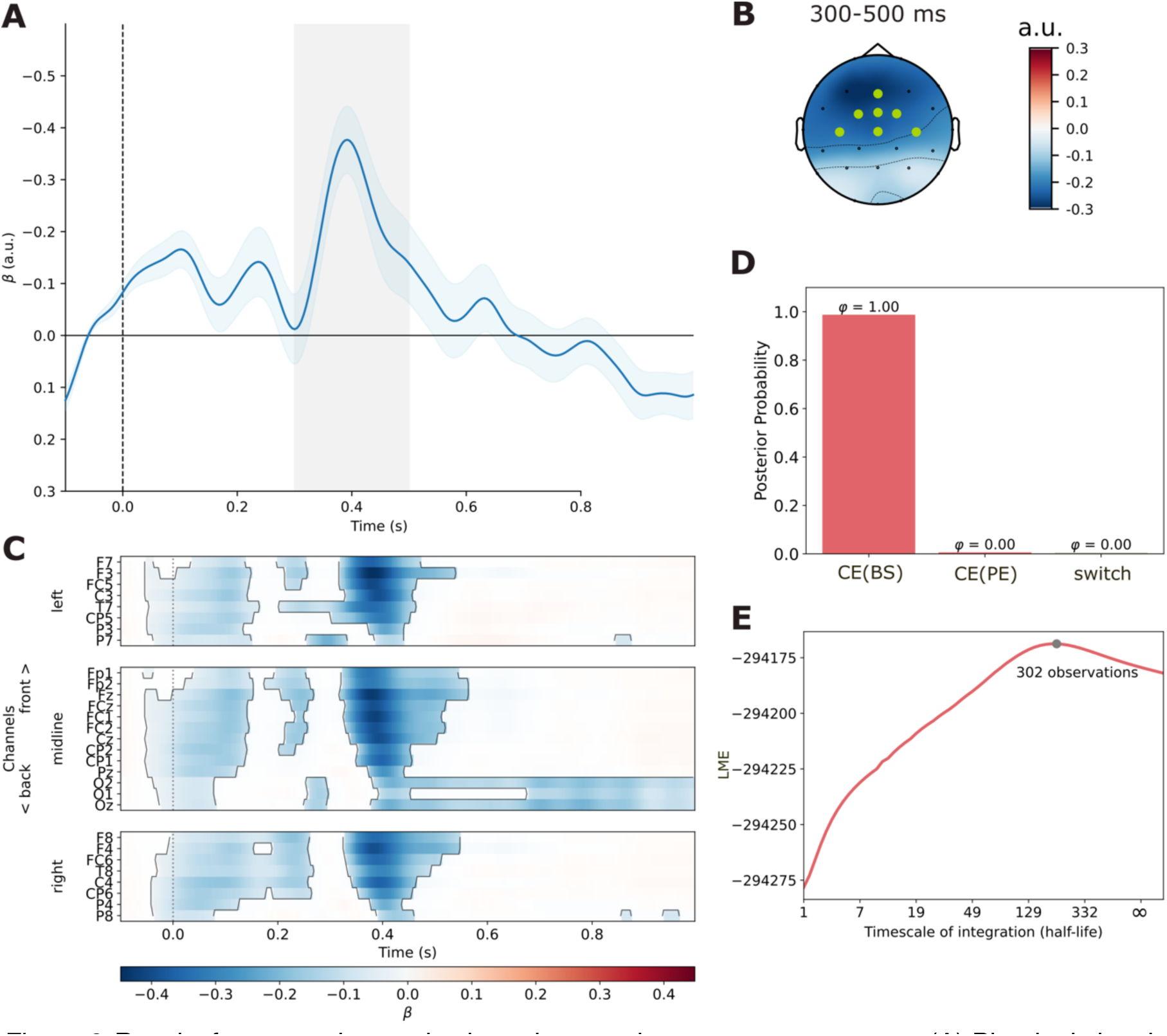
Results for semantic surprise based on previous category exposure. (A) Plot depicting the mean *β* values (grand-average rERP) of the category exposure model, with tau parameters optimized for the time window shaded in gray. (B) Topographical distribution for the time window shaded in gray (300-500 ms) with the ROI for the rERP plot marked by green dots. (C) Plot showing significant effects of category frequency according to the TFCE analysis. (D) Results from a Bayesian model comparison. Illustrated are posterior probability and protected exceedance probability for the category exposure model and two measures of surprise (BS and PE) as well as the non-probabilistic model representing category switches only. (E) Average log model evidence (LME) values corresponding to the parameter *τ*, which determined the time window of integration, here plotted as stimulus half-life.

**Figure 7.**
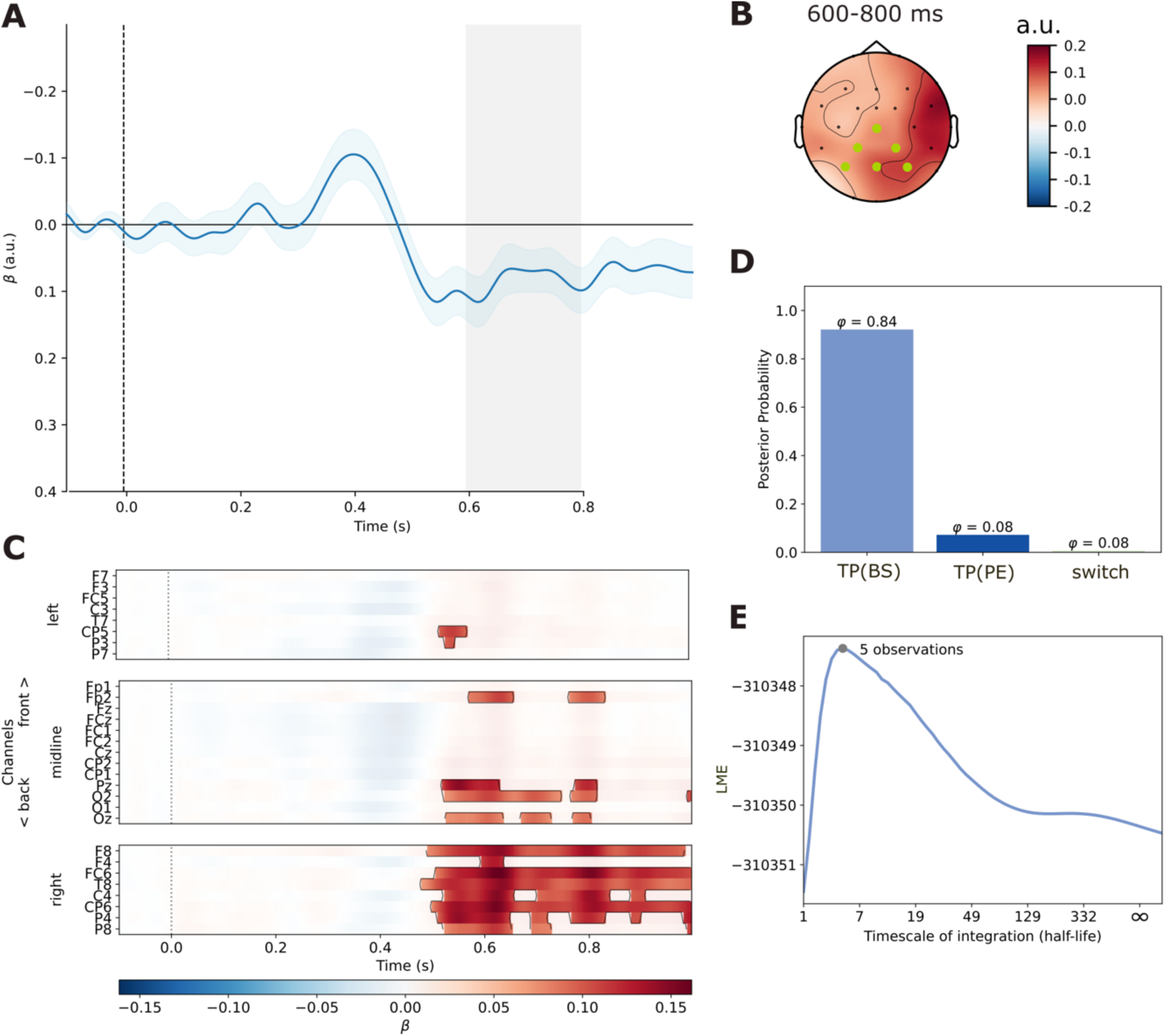
Results for semantic surprise based on the categories’ transition probabilities. (A) Plot depicting the mean *β* values (grand-average rERP) of the transition probability model, with tau parameters optimized for the time window shaded in gray. (B) Topographical distribution for the time window shaded in gray (600-800 ms) with the ROI for the rERP plot marked by green dots. (C) Plot showing significant effects of transition probability according to the TFCE analysis. (D) Results from a Bayesian model comparison. Illustrated are posterior probability and protected exceedance probability for the transition probability model and two measures of surprise (BS and PE) as well as the non-probabilistic model representing category switches only. (E) Average log model evidence (LME) values corresponding to the parameter *τ*, which determined the time window of integration, here plotted as stimulus half-life.

Single-trial brain responses before 500 ms after stimulus onset in the main sequence were primarily modulated by primarily modulated by semantic surprise based on previous exposure to the categories. This modulation is especially strong in the 300-500 ms time window and shows a similar spatial signature (Figure 6) as the mismatch N400 ERP effect (difference between standards and deviants: Figure 5). This quantitative characterization of the N400 is generally in line with the literature, as the ERP has been suggested to reflect unpredicted semantic information based on experimental (e.g., Federmeier et al., 2007; Kuperberg et al., 2020), verbally descriptive (e.g., Bornkessel-Schlesewsky & Schlesewsky, 2019), and computational work (Lindborg et al., 2023; Rabovsky et al., 2018; Rabovsky & McRae, 2014). In the current paradigm, the estimation of probabilities based on previous exposure seems to happen over a large timescale of integration. The best CE model therefore didn’t reflect local probabilistic semantic priming, as initially hypothesized based on Lindborg et al. (2023). Instead, N400 amplitudes in the current paradigm were influenced by repetition suppression and a general pre-activation of all semantic categories. This led to a reduction of amplitudes with time, as over the course of the experiment incoming items eventually have very little new semantic information (none of the semantic categories are surprising) and elicit only a small N400 amplitude.

The long timescale of integration seems to contradict previous modeling work specifically on the N400 ERP component by Lindborg et al. (2023), who modeled the component using Bayesian surprise in the same category exposure model. Specifically, they demonstrated that a model with a very local timescale of integration (*τ*=3; half-life ∼2)^1^ predicted N400 amplitudes. This conflict can be resolved by viewing the timescale of integration as a parameter that is adapted to the environment or task (Ossmy et al., 2013). In Lindborg et al.’s experiment participants were also presented with an oddball-like roving paradigm but spanning 10 different semantic categories (instead of 5 in the current study) and with trains per category consisting of more items than in the study reported here (mean of 6 items compared to 3). In a model estimating category probability based on very recent exposure (local integration), the estimate for the categories reflects a steadily increasing probability for nouns from the same category (i.e., its probability estimate becomes larger), and amplitudes increase with new semantic information at a deviant (low probability of other categories due to forgetting), describing the overall semantic mismatch effect (i.e., a local deviant compared to a standard) as an experience-based probabilistic process. In contrast, the reduced number of categories and shorter within-category trains of stimuli used in the current experiment has likely enabled the brain to form a probability representation across categories that, since global exposure frequencies were equal, led to repetition suppression for all categories and diminished the semantic mismatch response across time. In other words, all categories and concepts used in the current study were probably activated to a very high degree, leading to a floor effect on N400 amplitudes. Therefore, one explanation for the differences could be memory constraints interacting with the differences in paradigm (see also section “Implications for language related ERPs” below). We also want to emphasize that while there were effects of *local* exposure frequency on the N400 amplitude in Lindborg et al. (2023), neither experiment experimentally manipulated the *global* exposure frequency of semantic categories.

Interestingly, the local exposure model in Lindborg et al. (2023) also seemed to encode specifically the N400 ERP whereas the global integration over longer timescales in this analysis modulated earlier responses in addition to the N400. These early responses are oftentimes interpreted as word form prediction (for a comprehensive critical review see Nieuwland, 2019). Since it is highly unlikely that participants formed a prediction for the upcoming lexical item (their presentation was completely random), these early effects most likely reflect facilitated processing of lower-level features as the experiment progressed due to the massive repetition of items with time, even though this lower-level surprise was not modeled by the sequential learner, which instead focused on semantic categories.

### 4.2. Bayesian surprise based on transition probability modulates late brain responses

After the presentation of the main sequence in which global transition probabilities could be learned, the ERP semantic mismatch response in the test sequence did not reflect these manipulations, meaning there was no difference between HTP and LTP deviants. Effects of global transition probabilities have previously been demonstrated in many language learning paradigms (for a review see Saffran & Kirkham, 2018) and in the auditory system (e.g., Koelsch et al., 2016). Even though this comparatively subtle manipulation showed no significant effect in the current study, ERPs revealed a graded response based on the semantic category of the previous trial: more surprising transitions (low to high: within-category, between-category, novel transition) seemed to correspond to a more positive P600 for the deviant. While the semantic mismatch response (within vs between-category transitions, which is equal to standard vs deviant) was diminished with repetition in the N400 time window, it didn’t seem influenced by repetition suppression in the P600 time window. The more positive response to deviants in novel transitions compared to known between-category transitions suggests that this effect is likely more than a deterministic mismatch response.

Bayesian sequential learners revealed that brain responses after 500 ms post stimulus onset were modulated by Bayesian surprise in a model learning the transition probabilities between semantic categories. More specifically, larger Bayesian surprise at this level seems to correspond to more positive late ERPs. This modulation shows a similar spatial signature (Figure 6) as the P600 mismatch response for the respective data (Figure 4).

Interestingly, the short timescale of integration and the respective number of observations influencing the neural responses (here: *τ*=7; half-life ∼5) that was found for the transition probability model is consistent with the number of items that can be held in working memory (i.e., 4 according to Cowan, 2001 or 7 according to Miller, 1956). Overall, these results indicate that in a semantic sequence, the brain can use the current semantic category to predict the transition probability to the next semantic category.

The role of transition probabilities as implemented in this abstract sequence paradigm is of course just a very basic approximation to the complex dependencies underlying language processing, especially at the level of meaning. Therefore, the results are not supposed to propose these transition probabilities specifically as a computation for predictive processing in natural language but can rather be seen as proof of concept that conditional statistics at the level of meaning can be inferred and influence the neural response. In the case of the current paradigm, transition probabilities between semantic categories were the most complex but useful statistic to be tracked by the brain.

The sensitivity of the P600 to conditional expectancy violations, such as transition probabilities, is reminiscent of another famous late positive-going waveform: The P300 is a widely studied neurobiological marker of regularity violations, such as a deviant stimulus following a sequence of standard stimuli or violations of global patterns (Bekinschtein et al., 2009; Squires et al., 1976; Strauss et al., 2015; Sutton et al., 1965; Wacongne et al., 2011). Interestingly, recent literature employing similar paradigms and learner models as the ones described here, found surprise in a transition probability model to modulate the neural response around the P300 time window even when controlling for frequency of exposure to the stimuli (Maheu et al., 2019; Pesnot Lerousseau & Schön, 2021). Late positive responses have been suggested to reflect a more general need for revising the model of the environment, with the latency depending on stimulus complexity (Donchin & Coles, 1988). Since the oddball-like roving paradigm presented here manipulated semantic categories, stimuli should require more processing time than in standard oddball paradigms outside the domain of language processing (Kutas et al., 1977). The reported sensitivity of late positive neural responses to transition probabilities in a semantic sequence is therefore also in line with proposals that the P600 and P300 might share a common underlying cognitive mechanism (Bornkessel-Schlesewsky et al., 2011; Contier et al., in press; Coulson, 1998; Münte et al., 1998; Sassenhagen & Fiebach, 2019; van de Meerendonk et al., 2010). The short time window of integration found for the estimation of transition probabilities could also be explained by a conscious search for patterns (Bekinschtein et al., 2009; Dehaene et al., 2015; Wang et al., 2015). Since the statistics in the semantic sequence were not task-relevant and none of the participants reported to have noticed or searched a global pattern when asked after the experiment, we attribute only a minor role to controlled search processes. An explicit manipulation of task relevance and a replication of the paradigm with an active prediction or learning task would be an exciting future perspective, especially given the absent effects of the global transition probability pattern (HTP vs. LTP) in the test phase and the sensitivity of both the P600 and P300 to task demands and cognitive control (Brothers et al., 2022; Contier et al., 2022; Gratton et al., 2018; Leckey & Federmeier, 2020; Xu et al., 2021).

### 4.3. Implications for language related ERPs

The primary influence on N400 amplitudes in the current experiment was Bayesian surprise based on previous exposure during a relatively long period. This most likely reflects continuous activation of all five involved semantic categories due to massive repetition. This becomes apparent when considering that, across participants, semantic categories of on average 302 previous words seem to modulate N400 amplitudes to a substantial degree, which is equivalent to a steady activation of all semantic categories throughout the experiment.

Influences of transition probability surprise on the N400 could only be obtained in a post-hoc test focusing specifically on the N400 time window in a centro-parietal ROI rather than in our standard TFCE analysis spanning all electrodes and time points and controlling for multiple comparisons. While these results need to be treated with reservation until replicated, it seems plausible to assume that the continuous activation induced via massive repetition in the current paradigm created some sort of floor effect for N400 amplitudes (very small N400 amplitudes for all stimuli), making it difficult to corroborate more subtle influences. As noted above, in a similar paradigm with less repetition due to more different semantic categories (10 compared to 5 here) and longer trains of stimuli corresponding to the same category (mean of 6 items as compared to 3 here), N400 amplitudes were modulated by Bayesian surprise based on just very recent exposure, basically reflecting probabilistic semantic priming effects (Lindborg et al., 2023). Given that the current paradigm with its massive repetitions diminished even these probabilistic semantic priming effects in favor of long-term continuous repetition suppression, it may not be too surprising that potential influences of transition probability surprise on the N400 were hard to substantiate. Given ample evidence from sentence processing showing clear evidence for context-based predictability effects (Coulson et al., 2005; Kutas, 1993; Kutas & Hillyard, 1980, 1984; Van Petten, 1993), which constitute a form of complex semantic transition probability effects, and the positive evidence in our post-hoc test, we thus tentatively conclude that N400 amplitudes could be influenced by Bayesian surprise based on semantic transition probabilities beyond recent exposure, but that massive stimulus repetition can substantially diminish these influences (similar to the diminished semantic mismatch response in the test phase). Future work should replicate these influences in a paradigm with fewer repetitions.

On the other hand, transition probability effects on P600 amplitudes were strong despite the massive repetition (again, similar to the semantic mismatch response in the test phase). This is interesting in that it fits the idea that N400 amplitudes may reflect activation update in semantic memory where an already activated representation cannot increase its activation much further, while P600 amplitudes may reflect update of representations in working memory (see Rabovsky et al., 2018; Rabovsky & McClelland, 2020, for discussion) via interactions between frontal regions responsible for attention and control and temporal regions responsible for semantic representations. Crucially, only the semantic representations would be continuously active due to repetition. Linking P600 amplitudes to working memory update also seems to fit the short timescale of integration for transition probability surprise and the correspondingly small number of previous stimuli influencing the neural responses (*τ*=7; half-life ∼5). This would also be in line with the above-discussed link between the P600 and P300 components as the P300 has also been linked to context update in working memory (e.g., Polich, 2007) when the learner’s model of the environment must be revised (e.g., Donchin & Coles, 1988). Transition probability effects on the P600 in our semantic oddball paradigm seem to speak against the idea that the P600 reflects a very specific linguistic level of processing such as semantic integration in a sentence including thematic role assignment (Brouwer et al., 2017, 2021). Rather, the P600 seems to reflect Bayesian surprise and update based on conditional probabilities at a relatively high level of complexity (in this case semantic transition probabilities, but often also based on syntactic or pragmatic factors) independent of the specific linguistic structure of the input and the to-be-inferred representation.

### 4.4. Conclusion

Predictive processing as a concept spans multiple models and theories, but the essential prerequisite is the ability to infer a predictive pattern from the environment. Here we showed that such statistical inference is possible in a semantic sequence. Using Bayesian sequential learner models and EEG, we found that the brain can infer the hidden statistics in the sequential semantic input and continuously adapts its probability estimates for semantic categories.

Intriguingly, the results suggest that statistics of varying complexity might be used by the brain in semantic processing. At least in highly abstracted tasks including massive repetition such as the one employed here, components such as the N400 (as well as earlier components) seem to be best explained by simpler statistics such as probability estimates based on previous exposure that can model repetition suppression over the course of the experiment. Late positive deflections (corresponding to the P600 component) denoted a sensitivity to violated expectations based on conditional predictions, specifically transition probabilities calculated on a local timescale. This difference might be due to massive repetition having a strong impact on the activation and update of perceptual and semantic representations as reflected in early and mid-latency brain responses including the N400 while working memory update as presumably reflected by the P600 was less influenced by repetition. Global manipulations of transition probability were not reflected in the ERP response after the main sequence in which they could have been learned. Importantly, local transition probability inference as modeled here is most likely just a minimal statistical approximation of what is underlying the complex task of predictive language processing. However, the simplified paradigm allowed us to compare the findings to other statistical learning results and touch on the question of domain-specificity, as the current results revealed a striking similarity to results outside language processing. Together, the results are a step towards understanding not only the consequences of violated linguistic predictions, which is the focus of traditional ERP literature but also the continuous trial-by-trial adaptation of these predictions during language processing.

## Supplementary material

**Figure S1.**
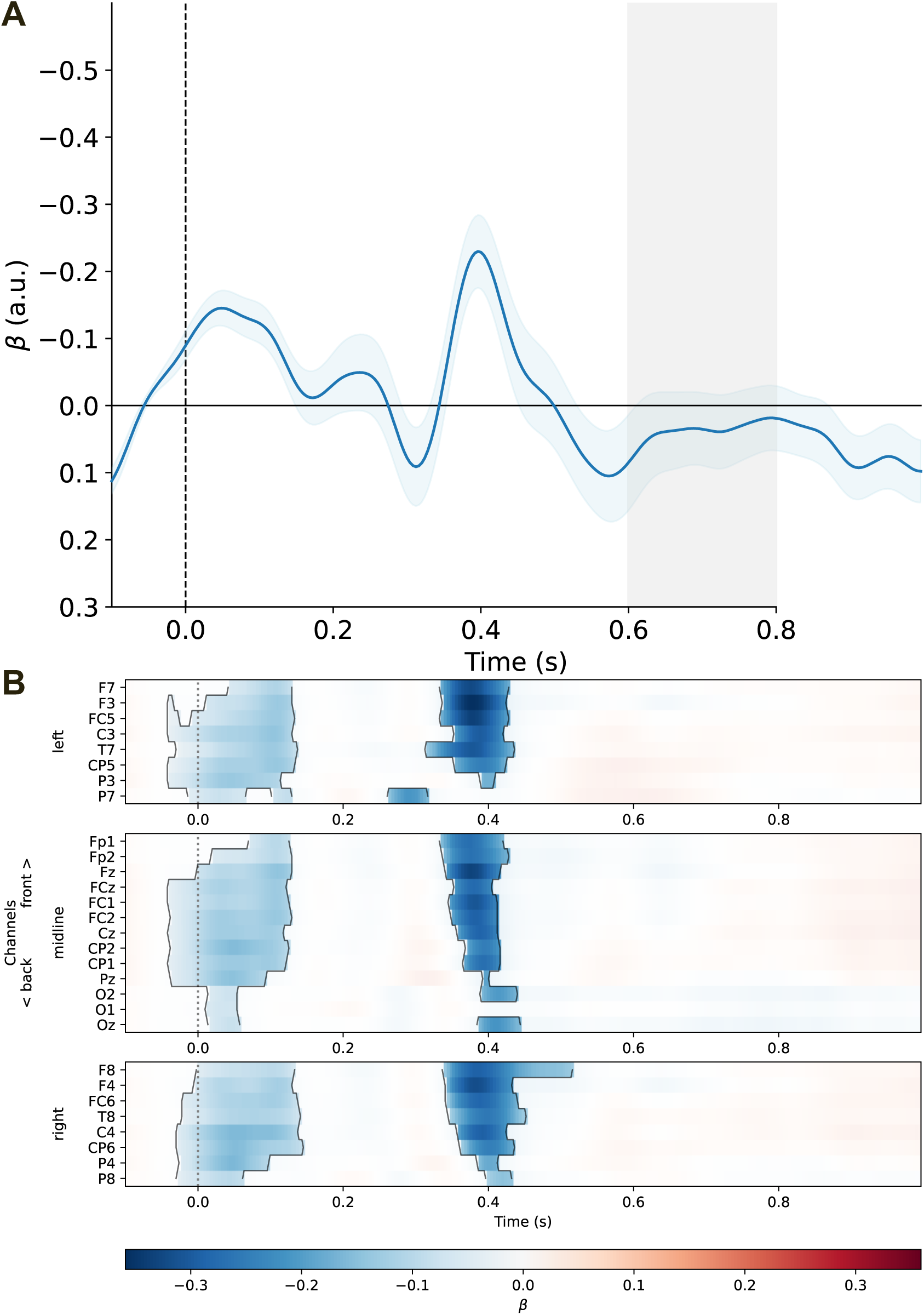
Category exposure model results. (A) Plot depicting the mean beta values (grand-average rERP) of the category exposure model, with tau parameters optimized for the time window shaded in gray. (B) Plot showing significant effects of category exposure according to the TFCE analysis. Model optimized for the 600-800 ms time window.

## Data availability

Modeling code, ERP analysis script, and averaged data are available on OSF: https://osf.io/jaf5s/. The raw EEG data cannot be made available in a public repository due to the privacy policies for human biometric data according to the European General Data Protection Regulation (GDPR) but can be requested from the corresponding other upon reasonable request.

## Author contributions

Conceptualization: AH, AL, MR; Data curation: AH; Formal analysis: AH; Funding acquisition: MR; Methodology: AH, AL; Software: AH, AL; Supervision: MR; Visualization: AH; Writing –original draft: AH, MR; Writing –review & editing: AH, AL, MR

## Acknowledgments

We thank Julia Kirschner and Dorottya Czárán for their help with data collection. We would also like to thank the HPC Service of ZIM, Universität Potsdam, for computing time.

---

**Figure S2.**
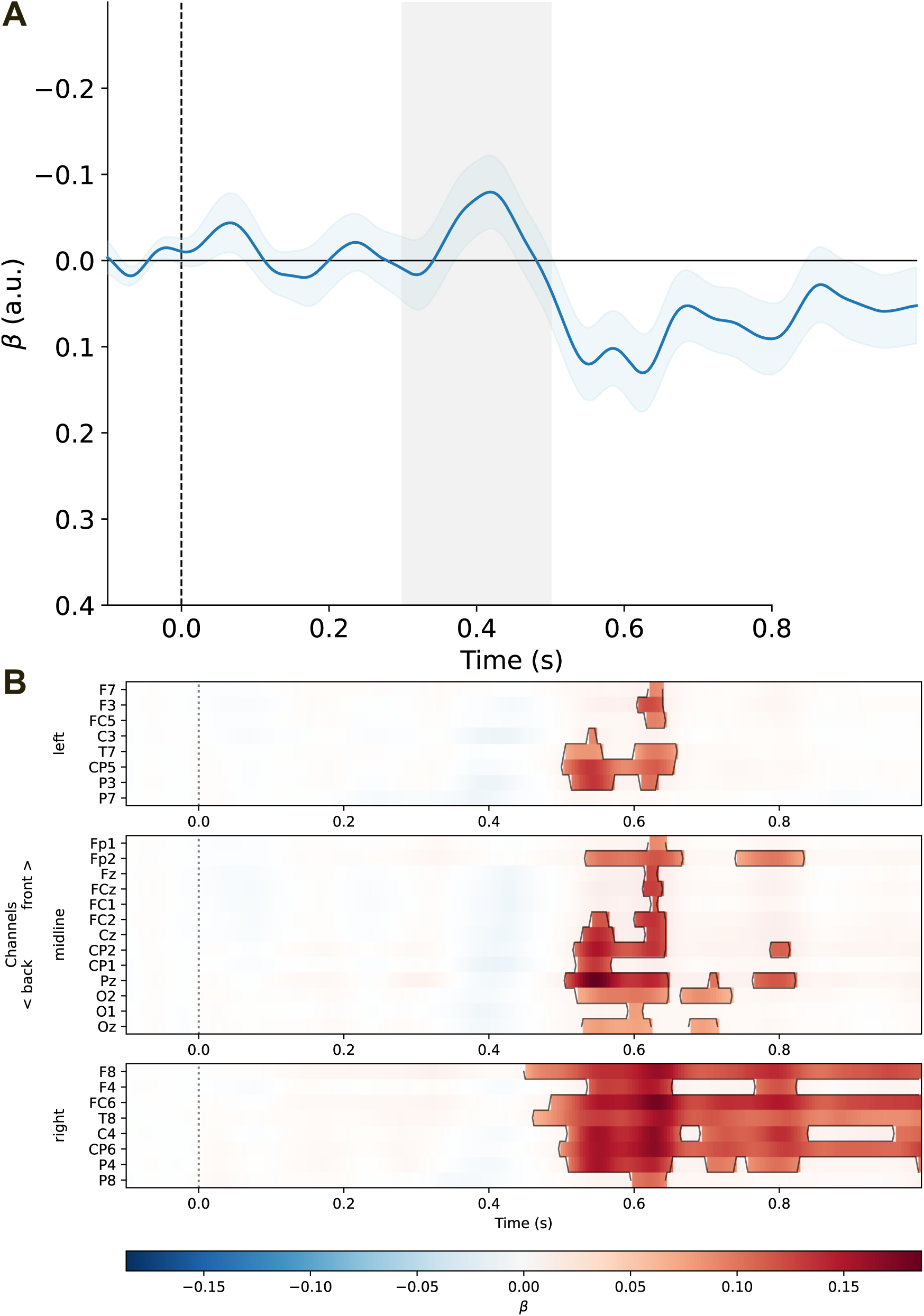
Transition probability model results. (A) Plot depicting the mean beta values (grand-average rERP) of the transition probability model, with tau parameters optimized for the time window shaded in gray. (B) Plot showing significant effects of transition probability according to the TFCE analysis. Model optimized for the 300-500 ms time window.

## Footnotes

1 This parameter was estimated across participants and not on a by-participant level as in the current paper. See also Gijsen et al. (2021), Grundei et al. (2023) and Maheu et al. (2019) for a similar by-participant approach and Musiolek et al. (2019) for findings from a semantic oddball task that indicate the necessity of by-participant parameters.

